# Multiple incursion pathways for *Helicoverpa armigera* in Brazil show its genetic diversity spreading in a connected world

**DOI:** 10.1101/762229

**Authors:** Jonas A. Arnemann, Stephen H. Roxburgh, Tom Walsh, Jerson V.C. Guedes, Karl H.J. Gordon, Guy Smagghe, Wee Tek Tay

## Abstract

The Old World cotton bollworm *Helicoverpa armigera* was first detected in Brazil with subsequent reports from Paraguay, Argentina, Bolivia, and Uruguay. This pattern suggests that the *H. armigera* spread across the South American continent following incursions into northern/central Brazil, however, this hypothesis has not been tested. Here we compare northern and central Brazilian *H. armigera* mtDNA COI haplotypes with those from southern Brazil, Uruguay, Argentina, and Paraguay. We infer spatial genetic and gene flow patterns of this dispersive pest in the agricultural landscape of South America. We show that the spatial distribution of *H. armigera* mtDNA haplotypes and its inferred gene flow patterns in the southwestern region of South America exhibited signatures inconsistent with a single incursion hypothesis. Simulations on spatial distribution patterns show that the detection of rare and/or the absence of dominant mtDNA haplotypes in southern *H. armigera* populations are inconsistent with genetic signatures observed in northern and central Brazil. Incursions of *H. armigera* into the New World are therefore likely to have involved independent events in northern/central Brazil, and southern Brazil/Uruguay-Argentina-Paraguay. This study demonstrates the significant biosecurity challenges facing the South American continent, and highlights alternate pathways for introductions of alien species into the New World.

## Introduction

Biological invasions are major ecological phenomena that influence the worldwide distribution of species. They can be a major driver of ecological change, affecting conservation (loss of biodiversity and species extinction [1, 2]), human health (e.g., the Zika virus transmitted by the invasive *Aedes aegypti* [3]) and agriculture (e.g., the spread of the cotton boll weevil, *Anthonomus grandis* through the Americas [4]; the introduction of fall army worm *Spodoptera frugiperda* into Africa from the New World [5]). In Brazil alone, 35 new pests have been detected [6] or confirmed [7, 8] in the last 10 years.

Invasive insects cost a minimum of US$70 billion per year globally for goods and services [9]. The Old World cotton bollworm, *Helicoverpa armigera* is considered a major agricultural pest with an estimated annual global cost to agriculture of over USD$5 billion [10], and Kriticos et al. [11] estimated that the arrival of *H. armigera* into North America would put at risk an extra USD$78 billion of agricultural output. A number of life history traits predispose *H. armigera* to be a highly successful insect pest [12]. These include: (i) high polyphagy, where larvae of this insect pest are known to feed on over 180 plant hosts from at least 68 plant families[13]; (ii) its long distance migratory ability with migrations of up to 2,000km [14–17]; (iii) the ability to enter a facultative diapause as pupae under unfavourable environmental conditions such as extreme high or low temperatures [12]; and (iv) high fecundity and a short generation time, capable of completing up to 10 to 11 generations per year [12,18,19]. Such fast generation times could aid in building population size, and thus contribute to a successful invasion [20].

In Brazil, *H. armigera* was confirmed in January-February 2013 [21, 22], and the incursion has resulted in over USD$800 million in losses and control costs since 2012 [6, 23, 24]. The presence of *H. armigera* was also reported in rapid succession in neighbouring western and south-western countries, including Paraguay in October 2013 [25, 26], Argentina in August-October 2013 [27], and in 2014/2015 in Uruguay [7, 28]. *H. armigera* was also confirmed in the Caribbean countries of Puerto Rico [29] and Dominican Republic [30,31] and on the mainland of United States of America [29]. The detection of *H. armigera* from southern Florida down to Argentina over just 3 years demonstrates the speed at which this species has established in the New World. It clearly has the ability to spread through a connected landscape and via an island hopping, stepping-stone dispersal model. The pathway of *H. armigera* into the South American continent may be linked with commodity movements involving importations of agricultural and horticultural products from multiple Old World destinations [31]. Whilst multiple origins of *H. armigera* in Brazil are likely, it remained unclear whether these founder individuals arrived as a single or as multiple event(s). By examining the genetic signature of the populations found in the South American continent using mtDNA markers, it may be possible to utilise spatial genetic signatures of the insect to infer frequencies of introductions.

It is almost certain that *H. armigera* had been present in South America for a period of time [e.g., 22, 32–34] prior to its first identification in Brazil [21], remaining undetected due to its close morphology with the New World sister species *H. zea*, and the difficulty of detecting invasive pests at the early stages of incursions [35]. It is unknown whether the *H. armigera* populations detected across South America arose from a single, or multiple, original introductions, and where these were. A detailed population genetic study is needed to test the hypothesis that there were multiple unrelated introduction events into South America, which would have significant implications for biosecurity preparedness for the South American continent and the potential reintroduction of novel adaptive ecotypes [36] into the Old World [33, 34]. To test this hypothesis, we undertook the present study, in which we show that gene flow and spatial distribution patterns of *H. armigera* mtDNA haplotypes support multiple introductions of the *H. armigera* into South America, with the incursion(s) in the southern regions likely independent from the northern/central Brazilian incursions.

## Results

### PCR amplification and sequence analysis

All specimens from the southern/south-western regions of South America were successfully sequenced for the mtDNA COI fragment (GenBank accession numbers MG230495-MG230526; KU255535-KU255543 from [7]) using the Noc-COI-F/R primer pairs. Sequence identity searches against the NCBI GenBank database confirmed that all suspected moths matched (i.e., 99-100% nucleotide identity) published *H. armigera* sequences, and did not contain premature stop codons.

Unique amino acid substitutions were detected in three haplotypes (i.e., Harm_BC47, Harm_BC42, Harm_BC43). For Harm_BC47, this involved an L/V change, and in Harm_BC42 a I/M change. Both substitutions involved amino acids with hydrophobic side chain. In Harm_BC43 the unique amino acid involved an A to G change where both amino acids belonged to the ‘small’ category. All remaining unique haplotypes (e.g., Harm_BC06, BC23, BC24. BC34, BC37, BC39, BC44, BC45, BC46) had nucleotide transition substitutions at 3^rd^ codon positions. The most significant unique non-Brazilian haplotypes detected were Harm_BC13, Harm_BC16, and Harm_BC17. All three unique haplotypes shared SNPs with other haplotypes, indicating that they did not have unexpected base changes. Furthermore, these haplotypes were also detected multiple times in separate sequencing efforts.

The range of genetic distances (i.e., measures of genetic divergence/degree of differentiation) of *H. armigera* within Asia (China, India and Pakistan) and within Australia were both 0.00 – 0.04%, while within Europe (Germany and unknown sites) and within South America (Brazil, Argentina, Uruguay and Paraguay) were both 0.00 - 0.02%. Estimates of evolutionary divergence between all *H. armigera* sequences from Australia, Asia, Europe and South America were therefore likewise low and ranged between 0.00 - 0.04%. Observed nucleotide diversity between countries/continents in *H. armigera* ranged from 0.0024 ± 0.0004 (s.e.) to 0.0040 ± 0.0006 (s.e.) (Table 1).

**Table 1:** Comparison of *Helicoverpa armigera* partial mtDNA COI gene nucleotide diversity (π ± s.e.) and haplotype diversity (h ± s.e.) between different countries/continents.

| Location | Nucleotide diversity ($\pi$) | Haplotype diversity ($h$) |
| --- | --- | --- |
| Asia | $0.00360 \pm 0.00034$ | $0.912 \pm 0.024$ |
| Europe | $0.00238 \pm 0.00040$ | $0.738 \pm 0.082$ |
| Australia | $0.00403 \pm 0.00057$ | $0.882 \pm 0.047$ |
| South America | $0.00244 \pm 0.00015$ | $0.769 \pm 0.018$ |
| Asia+Europe | $0.00314 \pm 0.00027$ | $0.862 \pm 0.032$ |
| Asia+Europe+Australia | $0.00380 \pm 0.00026$ | $0.894 \pm 0.024$ |

### Haplotypes

Haplotypes were collated from previous works [37, 38, 7] and from this study. A total of 47 haplotypes were identified from 314 individuals that consisted of 44 individuals from Asia, 18 from Australia, 28 from Europe and 226 from South America (Suppl. Table 3). Four most prevalent mtDNA COI haplotypes identified in this study were designated Harm_BC01, Harm_BC02, Harm_BC03, and Harm_BC04.

Brazil shares 5 mtDNA COI haplotypes with Asia (Harm_BC01, Harm_BC02, Harm_BC04, Harm_BC06 and Harm_BC07) and also 5 haplotypes with Europe (Harm_BC01, Harm_BC02, Harm_BC03, Harm_BC04 and Harm_BC06). Haplotypes Harm_BC01 and Harm_BC02 were present in all Brazilian states, and the Harm_BC01 haplotype was shared with all other locations, except Argentina, Paraguay and Uruguay, which had unique haplotypes (Harm_BC43, Harm_BC44, Harm_BC46 and Harm_BC47). The haplotype Harm_BC02 in Brazil was shared with Asia, Europe and Uruguay. Brazil also had 12 unique haplotypes (Harm_BC05, Harm_BC14, Harm_BC23, Harm_BC24, Harm_BC34, Harm_BC35, Harm_BC36, Harm_BC37, Harm_BC38, Harm_BC39, Harm_BC42 and Harm_BC45), and 80% of Brazilian individuals belonged to haplotypes Harm_BC01, Harm_BC02, Harm_BC03, Harm_BC04 and Harm_BC05.

### Helicoverpa armigera haplotypes distribution and validation of the null model

Results from validation of the null model indicated no tendency for the test to generate either Type I or Type II errors (i.e., having an almost perfect rectangular distribution of P-values across 10,000 statistical tests, with each test based on 10,000 random realisations of pseudo-observed data). At the 5% confidence level, almost exactly 5% of tests yielded statistically significant results with randomly generated data for the full dataset (501/10,000 = 0.0501 (Suppl. Fig. 1a); and when testing for Brazil vs. non-Brazil 496/10,000 = 0.0496 (Suppl. Fig. 1b)).

### Whole Table Analysis

Figure 1 shows the distribution of 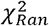 under the null model. No random realisations were found to have a value of the test statistic greater than or equal to that observed 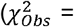 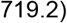, thereby indicating highly significant non-randomness within the observations (P = 0.000), and strong support that at least one haplotype/location observation has observations that are either more or less than expected by chance alone.

**Fig. 1:**
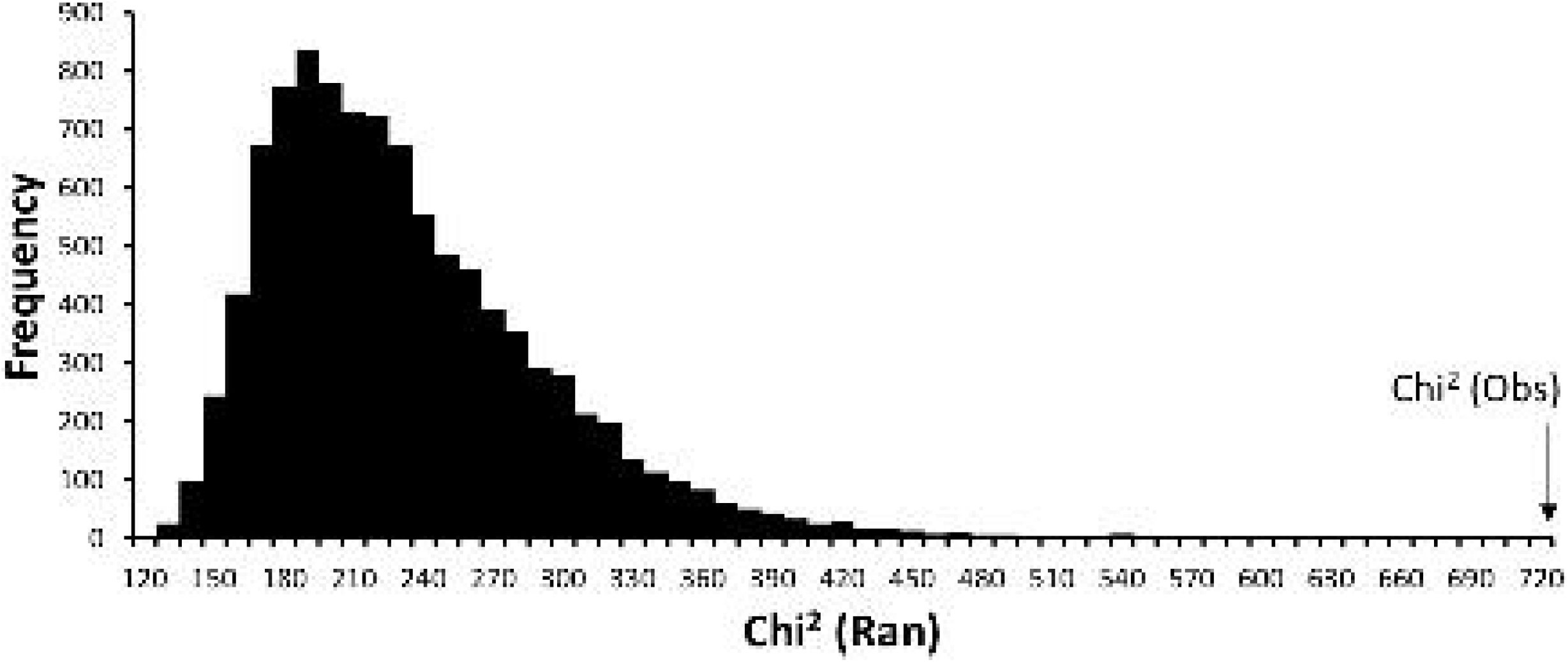
Distribution of 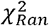 under the null model, and the location of 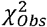

### Sub-table analysis

Evidence from above (Whole Table Analysis) strongly supported some mtDNA haplotypes were differentially distributed across sampling sites, and the *TS*_*DIFF*_ analysis was therefore used to further identify those haplotypes that were unduly rare or common across the locations. The results indicated haplotype Harm_BC01 and Harm_BC02 were both simultaneously under-represented in Argentina (ARG), Paraguay (PRY) and one Brazilian state (BA), and overly represented in another Brazilian state (PI) (Figure 2). The analysis also found evidence to support haplotypes Harm_BC03 and Harm_BC04 as being sporadically overly represented in three Brazilian states (BA, MT and RS) (dark blue cells, Figure 2).

**Figure 2.**
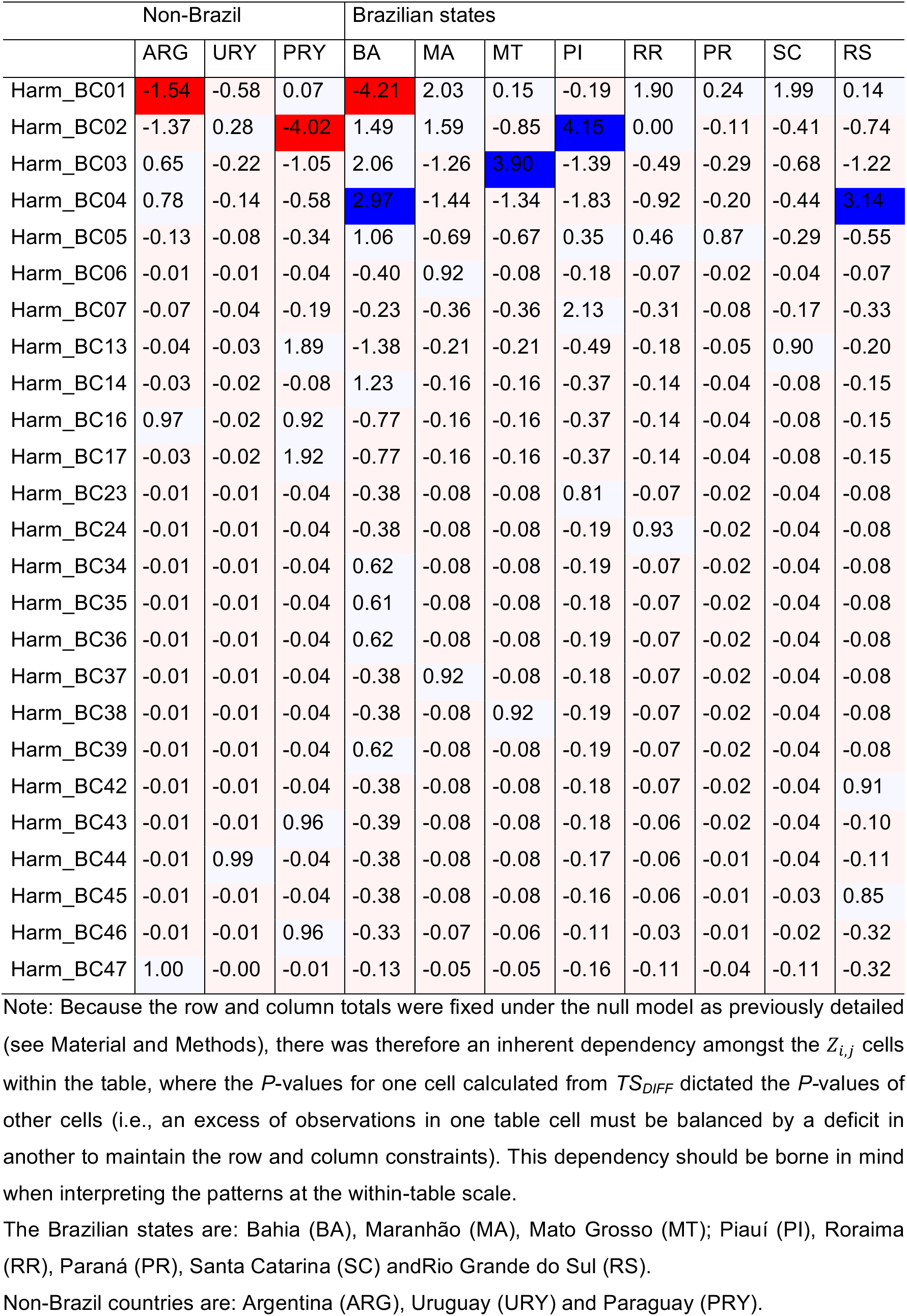
*TS*_*DIFF*_ values and indication of statistically significant deviations from the null model based of a false discovery rate (FDR) of 0.05 [67]. Pale red cells are cells with a non-significantly lower number of observations than expected at random, and pale blue cells non-significantly higher numbers of observations. Dark red cells indicate significantly lower numbers of observations. Dark blue cells indicate significantly higher numbers of observations

### Brazil vs. Non-Brazil

In the ‘Brazil vs. non-Brazil’ treatment of haplotype distribution data (Suppl. Table 4; Figure 3), non-randomness of haplotype distribution within the matrix was again confirmed by the 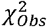 analysis (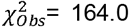, P-value < 0.000; Fig. 3). The irregular distribution of the test statistic in Figure 2 reflected a smaller dataset and, therefore, fewer possible combinations of allowable observations to fulfil the row and column constraints.

**Fig. 3:**
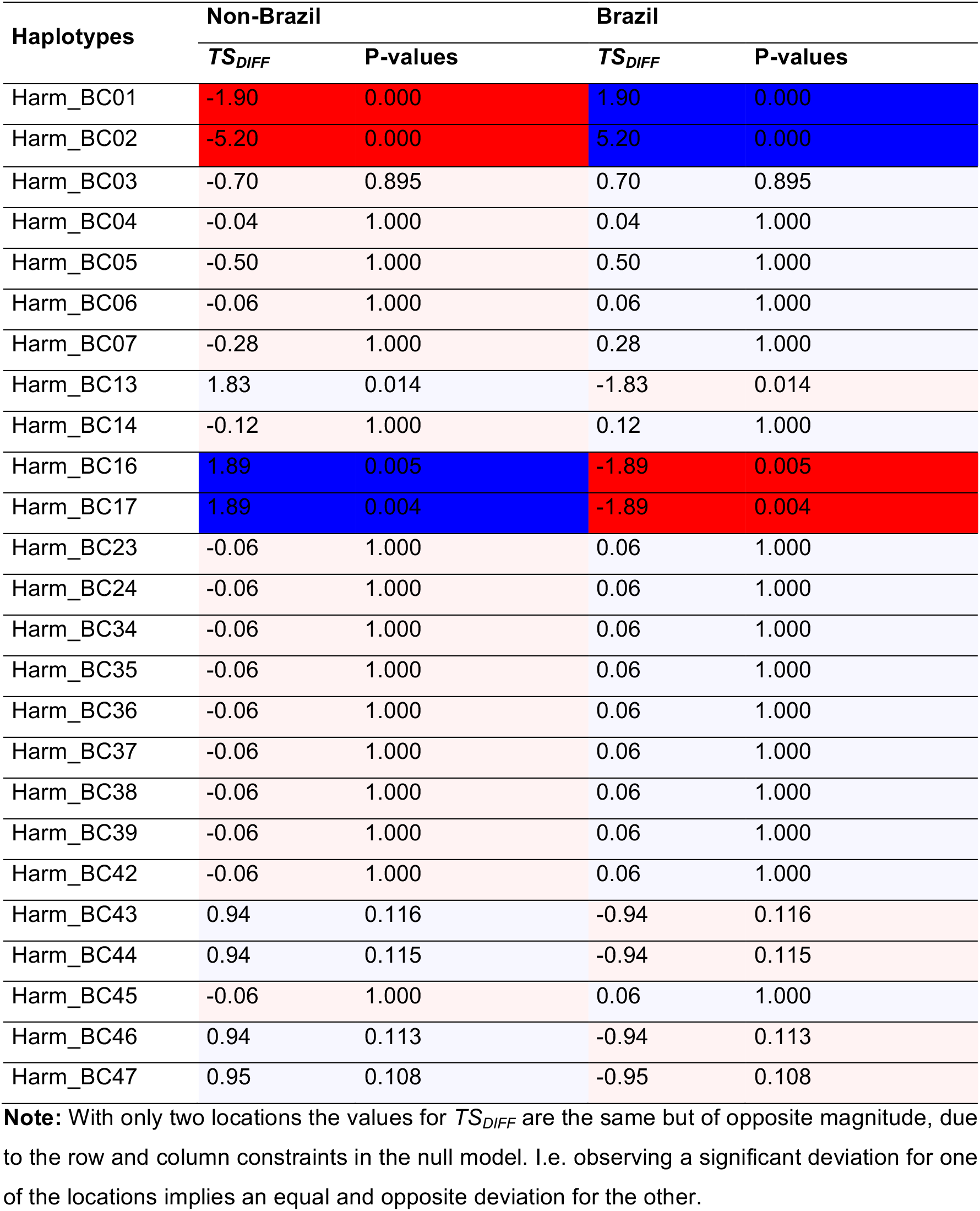
*TS*_*DIFF*_ for the Brazil vs Non-Brazil comparison of associated P-values

Consistent with the Whole Table Analysis (i.e., Figure 2), at the scale of the individual haplotypes, Harm_BC01 and Harm_BC02 were simultaneously under-represented in non-Brazil and over-represented in Brazil. For haplotypes Harm_BC13, Harm_BC16 and Harm_BC17 the opposite was true. A trend for over-representation of unique haplotypes Harm_BC43, Harm_BC44, Harm_BC46 and Harm_BC47 at Non-Brazil locations was also detected in this analysis, but deemed statistically insignificant under the Benjamin and Hochberg (1995) false discovery rate correction. The difference in the strength of statistical significance between the two analyses (Figure 2 and Figure 3) is due to a smaller allowable number of permutations when allocating observations at random to the two aggregated spatial categories.

### AMOVA and F_ST_ analysis

Overall *F*_ST_ estimates based on the partial mtDNA COI gene, when treating our data set as Brazilian, non-Brazilian (Uruguay, Argentina and Paraguay), and Old World samples, showed that a significant *F*_ST_ value (0.2879) was detected between Brazilian and non-Brazilian samples, indicating significant population sub-structure (i.e., low gene flow) between these populations (Suppl. Table 5). In contrast, the low *F*_ST_ estimate (0.0742, see also Tay et al. [31]) suggested consistent gene flow between Brazilian and Old World populations. Between the Non-Brazilian and the Old World populations, the significant pairwise *F*_ST_ estimate (0.2022) obtained suggested limited gene flow between these populations as inferred from the maternally inherited mtDNA COI gene.

At the fine-scale level, population pairwise *F*_ST_ (Suppl. Table 6) indicated limited gene flow between the Paraguay population and all of Brazilian populations, but not between the Paraguay, Argentina and Uruguay populations. Interestingly, limited gene flow also occurred between the Brazilian state of Rio Grande do Sul and the northern states of Piauí and Bahia, as well as with central Brazilian state of Mato Grosso. Similarly, the Argentinian population showed limited gene flow with Maranhão, Piauí and Bahia. Significant population substructure exhibited by Paraguay analysed with the whole Brazilian population would suggest the *H. armigera* population from Paraguay is unlikely to be recently established from Brazilian populations.

Analysis of molecular variance (AMOVA) among populations of *H. armigera* indicated that low levels of genetic variation existed between populations. Approximately 7% of variation detected could be attributed to that between individuals within groups, while the majority (>90%) of variation could be explained by the heterogeneous populations present across different geographic regions in the South American continent (Suppl. Table 7).

## Discussion

The rapid spread and establishment of *H. armigera* across much of the South American continent has generated a very large population with significant impacts on agricultural production. In contrast to usual invasions such as the African incursion of the New World fall armyworm *S. frugiperda* [16, 34, 39–42], the invasive *H. armigera* population in the New World appears to be very diverse. Furthermore, the spatial distribution of this diversity strongly suggests that the population has spread from two different regions of introduction. Statistical analyses of the haplotype distribution patterns show that some of the *H. armigera* haplotypes most commonly found in Brazil appeared to be uncommon in Argentina (i.e., Harm_BC01) and Paraguay (Harm_BC02), and conversely, that some of the less common and/or unique haplotypes found in the non-Brazilian countries (e.g., Harm_BC13, Harm_BC16, Harm_BC17) appeared disproportionately uncommon in Brazil. Furthermore, *F*_ST_ analyses suggest reduced gene flow between populations from the Cone Sul region (Southern Brazil, Argentina, Paraguay and Uruguay) and populations from either northern/central Brazil or from the Old World. Within individual countries, disproportionally over-represented haplotypes were identified (e.g., see Suppl. Table 2 and Suppl. Fig. 2), and within Brazil, the rare Harm_BC13 haplotype was detected in the State of Santa Catarina at sites located only approximately 350 km from Paraguay where this rare haplotype was also identified. Two other unique haplotypes (Harm_BC42, Harm_BC45) were also only identified in the southern state of Rio Grande do Sul in Brazil.

It is necessary to keep in mind that the northern and central Brazilian *H. armigera* may contain the same rare haplotypes as our southern region populations, although studies involving greater sample sizes from similar sampling periods [37, 38] did not detect these rare haplotypes. Nevertheless, rare haplotypes detected in Argentina, Paraguay and Uruguay have also not been detected in other Brazilian samples to-date. We were unable to include the mtDNA COI sequences of Tay et al. [31] due to different mtDNA COI regions being characterised. The study of Tay et al. [31] analysed substantial Brazilian populations from similar period and had identified similar haplotype frequency patterns as this study. Furthermore, Tay et al. [31] showed that within Brazil, the *H. armigera* population contained both globally common and globally rare haplotypes. At the national level, there were 13 rare haplotypes identified in Brazil reported, and is similar to the total number of 12 rare haplotypes identified in this study. Despite using different parts of the mtDNA COI gene, both studies have detected similar patterns of rare and common partial mtDNA COI haplotypes, suggesting that the results of our analyses are representative of the diversity present in Brazil at sampling time.

Population structure studies in *H. armigera* based on the mtDNA genes have found a general lack of substructure even for populations separated by considerable geographic distances (e.g. [51, 52]), and this finding is supported by studies based on limited nuclear markers (e.g., [15, 17, 53–57]). Anderson et al. [36] demonstrated differences between sub-species of *H. armigera* present in Australia/New Zealand (i.e., *H. armigera conferta*) and the Old World sub-species (i.e., *H. armigera armigera*), but not between global *H. armigera armigera* populations using genome-wide SNP markers. In northern/central Brazil, gene flow patterns of *H. armigera* showed non-significant levels of population substructure [24, 37] but exhibited reduced gene flow with southern South American populations of *H. armigera*. These southern South American populations, and particularly that of Paraguay and Argentina, also exhibited significant population substructure with Old World populations, suggesting that their origins differed from the origins of the founding populations present in northern/central Brazil.

Given that sufficient gene flow to prevent population structure had previously been detected in Brazilian *H. armigera* populations [24, 37, 38], and that *H. armigera* in Argentina, Paraguay and Uruguay [7, 25, 27] was confirmed shortly after the Brazilian detections, it would be a logical assumption that this likely represented natural migration, and/or movements of contaminated agricultural commodities from Brazil to the southern regions of South America. It was therefore unexpected to find unique mtDNA haplotypes in the southern South American populations that were as yet unreported in Brazil. Equally as unexpected was the lack of the most common haplotypes (e.g., Harm_BC01, Harm_BC02) in these countries given that they represented the major Brazil, and in fact, global, haplotypes.

To further explain the observed heterogeneous haplotype patterns across Paraguay, Uruguay, Argentina, and the southern Brazilian states of Rio Grande do Sul, Santa Catarina and Paraná (i.e., the Cone Sul region), two hypotheses may be put forward: (I) intrinsic factors associated with new biological incursions (e.g., stochastic lineage sorting, survival/reproductive variability, etc.) in a new environment, and (II) independent incursion pathways of *H. armigera* into South America.

In Brazil where the incursion of *H. armigera* was first reported, stochastic lineage sorting of founding populations (i.e., hypothesis I) could lead to the observed heterogeneous haplotype distribution patterns. This could involve factors such as lower population density and variability at population introduction phase and/or the lag-phase, high variability of reproductive success rates (e.g., see Gaither et al. [63]), variable adaptation success rates (e.g., differential response to attacks by parasitoids and/or predation rates [64]), susceptibility to viral/bacterial/fungal pathogen attacks, climatic stress, etc. to the novel New World environments at the early incursion stages.

In Brazil, across a number of studies, unexpectedly high genetic diversity of *H. armigera* has been detected [22, 24, 31, 37] as represented by multiple maternal lineages (Suppl. Table 2 e.g., Suppl. Fig.2). This would likely indicate a complex incursion history of *H. armigera* into Brazil, and may be associated with agricultural commodity movements from Old World regions into Brazil [31]. For example, repeated incursions/releases at different Brazilian sites could function as population source increasing propagule pressure, raising and maintaining diversity which is important in sustaining an incipient population [65, 66, 69] (i.e. hypothesis II). These suggested scenarios involving the highly volatile and variable periods of an exotic organism’s biology contrast the scenarios offered by Leite et al. [37], where repeated bottleneck effects such as potentially associated with differential pest control/management strategies were deemed likely factors that underpinned the rapid population expansion signatures in both *H. armigera* and the New World endemic and closely related *H. zea* (but see [31, 52]). In fact, *H. zea* in the New World was hypothesised as the outcome of an earlier incursion and the subsequent divergence (*ca*. 1.5 million years ago) from its common ancestor with *H. armigera* [52, 70], and involved a founder population with limited genetic diversity [71]. The *H. zea* genome, sequenced prior to the recent arrival of *H. armigera* in the New World, showed no evidence for subsequent introgression with *H. armigera*, and no evidence for the gain of additional genes affecting host use, but rather for the loss of genes already present in *H. armigera* [60].

Repeated introductions and high propagule pressure are increasingly being recognised as important factors that underpin the establishment of an alien species [2, 69]. With repeated introduction events, the likelihood of diverse maternal lineages that ultimately contribute to propagule pressure is high. Together with lineage sorting and stochastic processes (e.g., demographic, environmental [66]) experienced by the invasive species in the new environment, sampling of the mtDNA COI gene and the construction of a haplotype network will likely appear similar to one of a rapid population expansion (i.e., a ‘star-shaped’ haplotype network). This scenario of a ‘star-shaped’ haplotype network, as detected in *H. armigera* populations in the South Americas (see Fig. 2 of [37]), differed fundamentally to that reported for *H. zea* (i.e., Fig. 1 of [52]; Fig. 2 of [37]). Multiple introductions of an invasive pest insect that resulted in a mtDNA genetic signature similar to a rapid population expansion signature, has also been previously reported in Brazil (e.g. the Asian citrus psyllid *Diaphorina citri*, see [72]).

With the migration and dispersal ability of *H. armigera* in mind, the high frequency (i.e., 68%) of the two most common Harm_BC01 and Harm_BC02 haplotypes in Brazil populations, and a lack of population structures in northern/central Brazil (e.g., [37]) and the rest of the world, it was perhaps unexpected to observe statistically significant spatial mtDNA COI haplotype patterns and *F*_ST_ estimates in the Cone Sul region. The significant over- and underrepresented haplotypes in the Cone Sul region suggest that this population likely originated from somewhere outside the extensively sampled areas of central and northern Brazilian populations (i.e., hypothesis II). For example, haplotypes Harm_BC13 and Harm_BC17 for Paraguay and Harm_BC44 and Harm_BC47 for Uruguay and Argentina, respectively, were over-represented in these countries and to a lesser extent, also the over-representation of haplotypes Harm_BC43, Harm_BC44, Harm_BC46 and Harm_BC47,although the P-values (0.100 – 0.114) lie outside of the range usually considered significant, thereby adding support that these maternal lineages likely originated from non-Brazil source populations (i.e., hypothesis II).

The detection of the *H. armigera* genetic spatial signatures identified in this work provide the first insights into potential transnational patterns of the incursion of this species into the New World. This reflects opportunities to improve biosecurity protocols relating to phytosanitary practices of agricultural and horticultural commodity movements in the Cone Sul region. Although the haplotype distribution pattern from Rio Grande do Sul was excluded by our spatial analysis, pairwise *F*_ST_ estimates indicated a limited gene flow of the Rio Grande do Sul population with populations from three Brazilian states (see Suppl. Table 5), and coupled with the statistically significant over-representation of unique haplotypes in the Cone Sul region, our results therefore added support to the hypothesis of multiple introduction pathways of *H. armigera* into South America.

Similarly, the presence of limited gene flow between Rio Grande do Sul and Argentina populations and the various Brazilian populations might suggest different degrees of admixture between *H. armigera* from non-Brazilian and Brazilian populations. This observation should be further explored and should consider utilising genome-wide SNP markers to increase detection efficiency of population admixture [58]. Although the low sample size (n=2) in Uruguay has prevented meaningful interpretations of gene flow patterns based on the mtDNA COI marker, future genome-wide SNP markers studies on these individuals may enable migration patterns and the introduction history to be interpreted with more confidence. Taken as a whole, these *F*_ST_ results suggest that non-Brazilian *H. armigera* populations, and particularly those from Paraguay, followed by Argentina, and to a small extend that of the Rio Grande do Sul population, likely represent populations that were wholly or partially derived from alternative incursions event(s) from that detected in northern/central Brazil.

Single locus markers have clear limitations, however recent studies from multiple mtDNA markers [31] and from genome-wide SNP markers [58] have also demonstrated multiple introductions from globally diverse *H. armigera* populations in the invasive Brazilian populations. Taken as a whole, these findings provided evidence to support multiple independent introductions of *H. armigera* into the South American continent over the effect of stochastic lineage sorting. Our finding is not without precedent, with the globally invasive hemipteran whitefly *Bemisia tabaci* MED species (i.e., the real *B. tabaci* [59]) also being shown to have been independently introduced into this southern region of South America [60, 61] in addition to an earlier introduction elsewhere in Brazil [62].

Findings that populations of *H. armigera* in the South American continent likely also involved multiple introduction pathways was fortuitous, because the pattern identified in this study is unlikely to be maintained over time as mixing with other populations in South America and further incursions are likely. That incursions of *H. armigera* from the Old World potentially involved multiple pathways will have significant implications to pest and resistance management strategies in the New World. For example, populations of *H. armigera* around the world have developed resistance to conventional pesticides (e.g., [31, 73–78], and the South American populations have also arrived at least with resistance to pyrethroids [77] but also potentially with various enhanced allelochemical detoxification traits. Increasing genetic diversity is a key factor that underpins increasing invasion success [79, 80]. While significant levels of genetic diversity now exist in Brazil, and propagule pressure has also therefore decreased, the genetic make-up of these populations could be further bolstered by likely unrelated source populations from other parts of the Old World and this will further complicate and challenge management strategies. As pointed out by De Barro et al. [81], measures to restrict the recruitment of additional genetic diversity should be maintained even after establishment and spread have occurred, so as to avoid increasing the genetic diversity of damaging invasive pests.

## Material and Methods

### Sample collection and DNA extraction

Suspected *H. armigera* adults were collected using delta traps baited with the female sexual pheromone Iscalure armigera^®^ (ISCA Tecnologias LTDA, Ijuí, RS, Brazil) randomly installed in soybean fields in the Brazilian states Rio Grande do Sul, Santa Catarina and Paraná, and also in Uruguay, Argentina and Paraguay, in the 2014/15 cropping season (specifically, early April 2014; Table 2). Preservation of specimens, gDNA extraction procedures and PCR amplification and sequencing of partial mitochondrial DNA COI gene were done following PCR conditions as detailed in Arnemann et al. [7] using the primers Noc-COI-F (5’- GCGAAAATGACTTTATTCAAC -3’) and Noc-COI-R (5’- CCAAAAAATCAAAATAAATGTTG -3’).

**Table 2:** Collection sites, dates, and mtDNA COI GenBank accession numbers of *Helicoverpa armigera* specimens from Brazil (BRA), Uruguay (URY), Argentina (ARG) and Paraguay (PRY). Brazilian States are Rio Grande do Sul (RS), Santa Catarina (SC) and Paraná (PR).

| Code/state | Geographical coordinates Lat/Lon | Sampling date | GenBank Accession. |
| --- | --- | --- | --- |
| ARG | -32.1963222; -61.716597222 | 09-Apr-2014 | MG230495-MG230498 |
| PRY | -25.25508055; -57.56711111 | 12-Apr-2014 | MG230499-MG230502 |
| URY | -34.904344444; -54.93699166 | 03-May-2014 | MG230525-MG230526 |
| BRA-RS | -29.72653611; -53.561161111 | 10-Apr-2014 | MG230506-MG230517 |
| BRA-SC | -26.462400; -53.5098694444 | 06-Apr-2014 | MG230518-MG230524 |
| BRA-PR | -24.1583361111; -49.821575 | 25-Mar-2014 | MG230503-MG230505 |

### Selection of published mtDNA COI haplotype dataset

For our analysis, the following criteria underpinned our selection of mtDNA COI from publicly available sources: (A) only include published sequences from studies that had undergone critical review processes to avoid inclusion of untested haplotypes; (B) the sequences must originate from samples collected at a similar sampling time period as our material, (C) the published sequences must match our characterised partial mtDNA COI gene region, and (D) the populations must include northern/central Brazil to enable spatial comparisons to our Southern populations. Based on these criteria, sequences from three studies were chosen [7, 37, 38], with the published data from Arnemann et al. [7] also here included as part of the southern populations of *H. armigera*, while the studies of Leite et al. [37] and Mastrangelo et al. [38] represented the most comprehensive population diversity surveys at the mtDNA COI region in Brazil and fulfilled all four criteria.

### Sequence analysis of partial mtDNA COI gene

The programs Pregap and Gap4 within the Staden sequence analysis package [54] were used for editing DNA trace files and to assemble sequence contigs (i.e., haplotypes). Assembled mtDNA COI haplotypes were checked for premature stop codons that may be indicative of pseudogenes. Categorisation of global mtDNA COI haplotypes at the 5’ gene region, (Suppl. Table 1) and estimates of evolutionary divergence between all *H. armigera* individuals (i.e., between South America *vs.* Europe *vs.* Asia *vs.* Australia; n=314) involved 548bp of the mtDNA COI partial gene in the final dataset. The evolutionary divergence between haplotypes was estimated using the maximum composite likelihood model [44], and included all (i.e., 1^st^ + 2^nd^ + 3^rd^) codon positions using MEGA6 [45]. Estimates of haplotype diversity (*h* ± SE) and nucleotide diversity (π ± SE) were carried out using the molecular evolution software package DNA sequence polymorphism (DnaSP) v. 5.10.01 [46]. Assessments for presence of PCR errors, or nuclear mitochondrial sequences (NUMTs) involved (1) examination for presence of premature stop codons in the sequenced partial mtDNA COI gene region, (2) ascertaining for conservation of amino acid substitution patterns where presence of unique amino acid changes were further assessed for conservation of biochemical properties and/or their molecule sizes, and (3) sequence characterisation of rare haplotypes as confirmed by multiple independent PCR and sequencing efforts.

### Analysis of Helicoverpa armigera haplotypes spatial distribution patterns

To better investigate the spatial distribution patterns of *H. armigera* haplotypes in the South American continent, a matrix table was prepared for the frequencies of the 25 *H. armigera* mtDNA COI haplotypes identified to-date from 11 South American locations (i.e., from Brazilian samples sites: Bahia (BA), Maranhão (MA), Mato Grosso (MT), Piauí (PI), Roraima (RR), Paraná (PR), Santa Catarina (SC) and Rio Grande do Sul (RS); and from Argentina, Uruguay, and Paraguay; Suppl. Table 2), prior to performing a contingency table analysis using the χ^2^ statistic to detect departures, as detailed below. The statistical test was based on the randomisation of haplotypes x locations generated according to an appropriate null model (see ‘null model’ below), similar to Gotelli’s [47] analysis of species co-occurrence data.

### Null and alternative hypotheses

Due to the perceived unevenness of mtDNA COI haplotype spatial patterns, a statistical test was applied to ascertain whether the diversity and frequencies of individual haplotypes had occurred independently and at random across South American sampling sites, or whether there was evidence for spatial segregation. For such a test the null hypothesis is therefore that mtDNA COI haplotypes are distributed randomly across the locations. The alternative hypothesis therefore considers at least one haplotype as being either more or less common, in at least one location, than expected due to chance alone (i.e. haplotypes were differentially distributed across locations).

### Test statistics used

Equation (1) below was used for the χ^2^ statistic to detect departures from randomness:

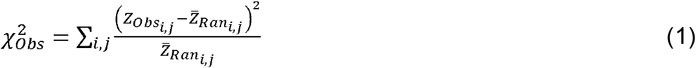

where 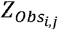 is the observation for row *i* and column *j* within the matrix (i.e. Suppl. Table 2), and 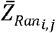 is the expected value for the same table entry calculated under the null hypothesis. If certain haplotypes and/or locations were disproportionately under or over-represented, then 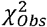 would be expected to fall within the extreme tail of the distribution of 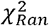 (i.e., the calculated value of the statistic when the null hypothesis is known to be true).

Although an analysis based on 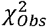 provides an overall test of haplotype randomness across the landscape, information on which location and/or haplotype combinations underpin any deviation from randomness will require an additional test statistic (i.e., equation (2)) to be applied at the scale of each haplotype/location combination:

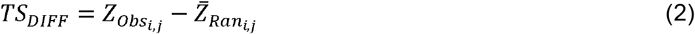

Equation (2) implies that when *TS*_*DIFF*_< 0.0 a haplotype would be less common than expected in location *j* due to chance alone, when *TS*_*DIFF*_> 1.0 the haplotype would be more common than expected in location *j* by chance, and when *TS_VAR_* = 0.0 the observed data conformed with the null model. Analysis of *TS*_*DIFF*_ can therefore be used to further explore primary circumstances (i.e., combinations of haplotype and location) and directionality (i.e., whether haplotypes were unexpectedly rare or common across the sites) if/when evidence of differentially distributed haplotypes across the landscapes was detected (i.e. to identify the genotypes that are unduly rare or common across the locations).

Determining the statistical significance of individual table entries is problematic because multiple comparisons are being simultaneously assessed. For example in Figure 2, there are 11 locations × 25 haplotypes = 275 cell-level tests, of which approximately 14 would be expected to show significance simply by chance (i.e. at the 0.05 level of confidence). To control for this ‘familywise’ error rate the method of Benjamini and Hochberg [67] was applied, using a false discovery rate of 0.10 [68]. This identified seven cell entries of interest in the full analysis (Figure 2), and five pairwise (Brazil /non-Brazil) comparisons of interest when data were pooled across locations (Figure 4).

**Fig. 4:**
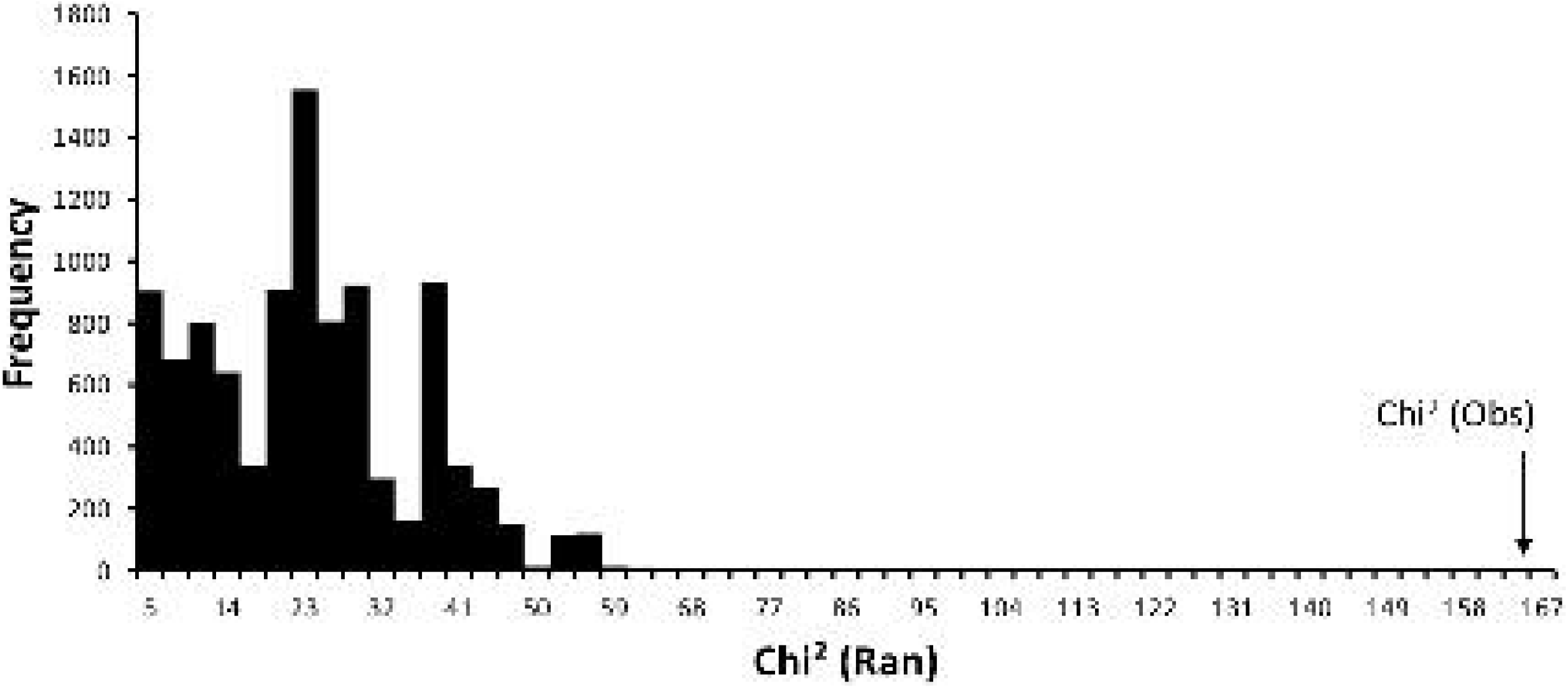
Distribution of 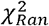 under the null model, and the location of 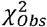 for the Brazil haplotypes vs non-Brazil haplotypes comparison

### Null model

The null model generated for this study involved distributing all 226 observations (i.e., the complete mtDNA COI dataset of *H. armigera* in South America; Suppl. Table 2) across haplotypes and locations at random, with the constraint of fixed row and column totals (i.e. the same number of observations of each haplotype, and the same number of observations for each location, are both retained in the randomised matrices). This treatment is necessary to ensure that the null model explicitly accounts for both unequal survey efforts at different sites and the overall lower frequencies of some haplotypes. The algorithm AS159 of Patefield [48] was used to generate random tables that constrained both row and column marginal totals.

This null model represents a non-parametric alternative to Fisher’s exact test, which is also based on fixed marginal totals, but which relies upon a chi-squared distribution to approximate the underlying exact distribution. This approximation can be poor if the data are sparse (see Suppl. Table 3), hence the decision to calculate *P-*values via random permutation.

### Calculating P-values

P-values for 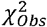 were calculated by enumerating the number of times 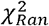 was less than or equal to 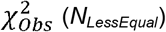, and also the number of times 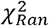 was greater than or equal to 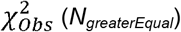, based on 10,000 randomly generated tables. The two-tailed P-value [49] is given by:

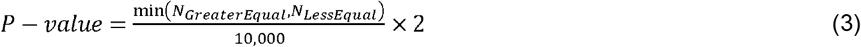

P-values for *TS*_*DIFF*_ were calculated similarly, but at the scale of individual table cells. Note that as the quantities *N*_*GreaterEqual*_ and *N*_*LessEqual*_can potentially overlap in their included values (i.e., both include an equality term), P-values can potentially exceed 1.0; in such cases values are rounded down to 1.0.

### Validating the null model

The statistical test was validated by creating *n* pseudo-observed data sets under the null model, followed by application of the test to each of the created data sets to confirm appropriate Type I and Type II error rates. At the 5% confidence level only 5% of pseudo random data sets should be statistically significant, and the expected P-value distribution across the *n* tests should be rectangular. If it is found that more than 5% of test results are significant at the 5% confidence level, then this indicates an elevated Type I error rate (i.e., a chance that the null hypothesis is incorrectly rejected when no real differences are actually present). Conversely, when less than the nominal number of pseudo random data sets yield a significant result it can lead to incorrect acceptance of the null hypothesis (i.e., concluding no significant difference even when one is actually present; (i.e., an elevated Type II error rate)).

The null model was validated through repeated analyses (10,000 times) using random pseudo-data reconstructed to match the n = 266 observations (Suppl. Table 2), but allowing observations to be randomly allocated to the matrices, and thereby ensuring conformity with the null hypothesis. Tables that had no observations for a given row or column during the constructing of pseudo-data were excluded. The code, the program and test data are provided as supplementary material (Suppl. Material 1).

### Brazil vs. Non-Brazil

The above analyses were repeated, but combining the location data to consider just two categories of samples – the Brazilian and the non-Brazilian (Argentina, Paraguay, Uruguay) samples. Note that this analysis was carried out as a guide to assist interpretation for the observed within-table patterns of deviations, and have not impacted our main findings which was the result from the full dataset (i.e., Figure 2) analysis.

### AMOVA and F_ST_ analysis

The population pairwise *F*_ST_ and AMOVA estimates were carried out using Arlequin 3.5.2.2 [39], and significance values were estimated with 10,000 permutations. For the AMOVA, populations were assigned into four groups to test molecular variation across geographical regions of Brazil (North: Roraima, Maranhão, Piauí; Central/Eastern: Bahia, Mato Grosso; South: Rio Grande do Sul, Santa Catarina, Paraná) and neighbouring countries (Argentina, Paraguay, Uruguay).

## Acknowledgements

We thank Gustavo Ugalde and Elgion Loreto for laboratory assistance. We thank Claudia C. C. Antúnez and Enrique Castiglioni for providing Paraguay and Uruguay samples, respectively. Clérison R. Perini, Cristiano De Carli, Glauber R. Sturmer, Luis E. Curioletti and Rubens A. Fiorin assisted with Brazilian *H. armigera* sampling and DNA preparation.

## Authors’ contributions

JAA, WTT, JVCG, KHJG, TW and GS conceived the ideas and designed methodology; JAA and WTT collected the data; JAA, WTT and SR analysed the data; WTT, TW, SR, KHJG and JAA led the writing of the manuscript. All authors contributed critically to the drafts and gave final approval for publication.

## Competing interests

We declare we have no competing interests.

## Funding

This study was funded by UFSM, CSIRO Health and Biosecurity ‘Genes of Biosecurity Significance’ (R-8681-1) and Ghent University.

## References

1. Lambertini, M. et al. Invasives: a major conservation threat. Science 333, 404–405 (2011).

2. Simberloff, D. et al. Impacts of biological invasions: what’s what and the way forward. Trends. Ecol. Evol. 28, 58–66 (2013).

3. Zanluca, C. et al. First report of autochthonous transmission of zikavirus in Brazil. Mem. Inst. Oswaldo Cruz 110, 569–572(2015).

4. Barbosa, S., Braga, S.R., Lukefahr, M.J. & Beingolea, G.O. Relatório sobre a ocorrência do bicudo do algodoeiro, Anthonomus grandis Boheman “Boll Weevil”, no Brasil e recomendações para a sua erradicação. Campina Grande: Embrapa CNPA (1983).

5. Goergen, G., Kumar, P.L., Sankung, S.B., Togola, A. & Tamò, M. First report of outbreaks of the fall armyworm *Spodoptera frugiperda* (J E Smith) (Lepidoptera, Noctuidae), a new alien invasive pest in west and central Africa. PLoS ONE 11, e0165632; 10.1371/journal.pone.0165632 (2016).

6. Lopes-da-Silva, M., Sanches, M.M., Stancioli, A.R., Alves, G. & Sugayama, R. The role of natural and human mediated pathways for invasive agricultural pests: a historical analysis of cases from Brazil. Agricultural Sciences 5, 634–646 (2014).

7. Arnemann, J.A. et al. Mitochondrial DNA COI characterization of *Helicoverpa armigera* (Lepidoptera: Noctuidae) from Paraguay and Uruguay. Genet. Mol. Res.15, 15028292; 10.4238/gmr.15028292 (2016).

8. Arnemann, J.A.et al. Soybean stem fly, *Melanagromyza sojae* (Diptera: Agromyzidae), in the new world: detection of high genetic diversity from soybean fields in Brazil. Genet. Mol. Res 15, 15028610; 10.4238/gmr.15028610 (2016).

9. Bradshaw, C.J.A. et al. Massive yet grossly underestimated global costs of invasive insects. Nat. Commun. 7, 12986; 10.1038/ncomms12986 (2016).

10. Sharma, H.C. Heliothis/Helicoverpa management: emerging trends and strategies for future research. New Delhi: Oxford& IBH Publishing Co. Pvt. Ltd. (2005).

11. Kriticos, D.J. et al. The potential distribution of invading *Helicoverpa armigera* in North America: is it just a matter of time? PLoS ONE 10, e0133224; 10.1371/journal.pone.0133224 (2015).

12. Fitt, G.P. Ecology of *Heliothis* species in relation to agroecosystems. Annu. Rev. Entomol. 34, 17–52 (1989).

13. Cunningham, J.P. & Zalucki, M.P. Understanding Heliothine (Lepidoptera: Heliothinae) pests: what is a host plant? J. Econ. Entomol. 107, 881–896 (2014).

14. Widmer, M.W. & Schofield, P. Heliothis: Dispersal and Migration. London: Tropical Development and Research Institute (1983).

15. Nibouche, S., Buès, R., Toubon, J.F. & Poitout, S. Allozyme polymorphism in the American cotton bollworm *Helicoverpa armigera* (Hübner) (Lepidoptera: Noctuidae): comparison of African and European populations. Heredity 80, 438–445 (1998).

16. Jones, C.M. et al. Genomewide transcriptional signatures of migratory flight activity in a globally invasive insect pest. Mol. Ecol. 24, 4901–4911 (2015).

17. Behere, G.T., Tay, W.T., Russell, D.A., Kranthi, K.R. & Batterham, P. Population genetic structure of the cotton bollworm *Helicoverpa armigera* (Hübner) (Lepidoptera: Noctuidae) in India as inferred from EPIC-PCR DNA markers. PLoS ONE 8, e53448; 10.1371/journal.pone.0053448 (2013).

18. Coaker, T.H. Investigations on *Heliothis armigera* in Uganda. B. Entomol. Res. 50, 487–506 (1959).

19. Raulston, J.R. et al. Ecological studies indicating the migration of *Heliothis zea, Spodoptera frugiperda* and *Heliothis virescens* from northeastern Mexico and Texas in *Insect Flight, Dispersal and Migration* (ed. Danthanarayana, W.) 128–144. Heidelberg, Germany: Springer-Verlag (1986).

20. Tayeh, A. et al. Biological invasion and biological control select for different life histories. Nat. Commun. 6, 7268; 10.1038/ncomms8268 (2015).

21. Czepak, C., Albernaz, K.C., Vivan, L.M., Guimaraes, H.O. & Carvalhais, T. Primeiro registro de ocorrência de *Helicoverpa armigera* (Hübner) (Lepidoptera: Noctuidae) no Brasil. Pesqui. Agropecu. Trop. 43, 110–113 (2013).

22. Tay, W.T. et al. A brave new world for an Old World pest: *Helicoverpa armigera* (Lepidoptera: Noctuidae) in Brazil. PLoS ONE 8, e80134; 10.1371/journal.pone.0080134 (2013).

23. Pomari-Fernandes, A., Freitas-Bueno, A. & Sosa-Gómez, D.R. *Helicoverpa armigera*: current status and future perspectives in Brazil. Current Agricultural Science and Technology 21, 1–7 (2015).

24. Mastrangelo, T. et al. Detection and genetic diversity of a heliothine invader (Lepidoptera: Noctuidae) from north and northeast of Brazil. J. Econ. Entomol. 107, 970–980 (2014).

25. Senave. Senave en alerta tras ingreso de peligrosa plaga agrícola. Available at http://www.abc.com.py/edicion-im7resa/economia/senave-en-alerta-tras-ingreso-de-peligrosa-plaga-agricola-629240.html (2013).

26. EPPO. EPPO Global Database. Available at: https://gd.eppo.int (2014).

27. Murúa, M.G. et al. First record of *Helicoverpa armigera* (Lepidoptera: Noctuidae) in Argentina. Fla. Entomol. 7, 854–856 (2014).

28. Castiglioni, E. et al. Primer registro de ocurrencia de *Helicoverpa armigera* (Hübner, 1808) (Lepidoptera: Noctuidae) en soja, en Uruguay. Agrociencia Uruguay 20, 31–35 (2016).

29. Gilligan, T.M., Goldstein, P.Z., Timm, A.E., Farris, R., Ledezma, L. & Cunningham, A.P. Identification of Heliothine (Lepidoptera: Noctuidae) Larvae Intercepted at U.S. Ports of Entry From the New World. J. Econ. Entomol. doi: 10.1093/jee/toy402. [Epub ahead of print] (2019).

30. EUROPHYT. European Union Notification System for Plant Health Interceptions Annual Report2014. Luxembourg: Food and Veterinary Office (2015).

31. Tay, W.T. et al. Mitochondrial DNA and trade data support multiple origins of *Helicoverpa armigera* (Lepidoptera, Noctuidae) in Brazil. Scientific Reports 7, 45302; 10.1038/srep45302 (2017).

32. Sosa-Gómez, D.R. et al. Timeline and geographical distribution of *Helicoverpa armigera* (Hübner) (Lepidoptera, Noctuidae: Heliothinae) in Brazil. Rev. Bras. entomol. 60(1); 10.1016/j.rbe.2015.09.008 (2016).

33. Jones, C.M., Lim, K.S., Chapman, J.W. & Bass, C. Genome-wide characterization of DNA methylation in an invasive lepidopteran pest, the cotton bollworm *Helicoverpa armigera*. G3: Genes, Genomes, Genetics 8(3), 779–787; 10.1534/g3.117.1112 (2018).

34. Tay W.T. & Gordon, k.H.J. (2019) Going global – Genomic insights into insect invasions. Curr. Opin. Insect Sci. https://doi.org/10.1016/j.cois.2018.12.002 (2019).

35. Crooks JS (2005) Lag times and exotic species: The ecology and management of biological invasions in slow-motion. Ecoscience 12(3), 316–329.

36. Anderson, C.J. et al. Hybridization and gene flow in the mega-pest lineage of moth, *Helicoverpa*. Proceedings of the National Academy of Sciences 115(19), 5034–5039; 10.1073/pnas.1718831115 (2018).

37. Leite, N.A., Alves-Pereira, A., Corrêa, A.S., Zucchi, M.I. & Omoto, C. Demographics and genetic variability of the new world bollworm (*Helicoverpa zea)* and the Old World bollworm (*Helicoverpa armigera)* in Brazil. PLoS ONE 9, e113286; 10.1371/journal.pone.0113286 (2014).

38. Mastrangelo, T. et al. Detection and genetic diversity of a heliothine invader (Lepidoptera: Noctuidae) from north and northeast of Brazil. J. Econ. Entomol. 107, 970–980 (2014).

39. Goergen, G., Kumar, P.L., Sankung, S.B., Togola, A. & Tamò, M. First report of outbreaks of the fall armyworm *Spodoptera frugiperda* (J. E. Smith) (Lepidoptera, Noctuidae), a new alien invasive pest in West and Central Africa. PLoS ONE 11(10), e0165632; 10.1371/journal.pone.0165632 (2016).

40. Cock, M.J.W., Beseh, P.K., Buddie, A.G., Cafá, G. & Crozier, J. Molecular methods to detect *Spodoptera frugiperda* in Ghana, and implications for monitoring the spread of invasive species in developing countries. Scientific Reports 7, 4103; 10.1038/s41598-017-04238-y (2017).

41. Nagoshi, R.N., Koffi, D., Agboka, K., Tounou, K.A., Banerjee, R. et al. Comparative molecular analyses of invasive fall armyworm in Togo reveal strong similarities to populations from the eastern United States and the Greater Antilles. PLoS ONE 12(7), e0181982; 10.1371/journal.pone.0181982 (2017).

42. Otim, M.H., Tay, W.T., Walsh, T.K., Kanyesigye, D., Adumo, S. et al. Detection of sister-species in invasive populations of the fall armyworm *Spodoptera frugiperda* (Lepidoptera: Noctuidae) from Uganda. PLoS ONE 13(4), e0194571; 10.1371/journal.pone.0194571 (2018).

43. Staden, R.B.K.F. & Bonfield, J.K. The Staden package, 1998. Methods in Molecular Biology 132, 115–130 (2000).

44. Tamura, K., Nei, M. & Kumar, S. Prospects for inferring very large phylogenies by using the neighbor-joining method. Proc. Natl. Acad. Sci. U S A 101,11030–11035 (2004).

45. Tamura, K., Stecher, G., Peterson, D., Filipski, A. & Kumar, S. MEGA6: Molecular Evolutionary Genetics Analysis version 6.0. Mol. Biol. Evol. 30, 2725–2729 (2013).

46. Librado, P. & Rozas, J. DnaSP v5: a software for comprehensive analysis of DNA polymorphism data. Bioinformatics 25, 1451–1452 (2009).

47. Gotelli, N.J. Null Model Analysis of species co-occurrence patterns. Ecology 81, 2606–2621 (2000).

48. Patefield, W.M. Algorithm AS 159: an efficient method of generating random *R*x*C* tables with given row and column totals. Applied Statistics 30, 91–97(1981).

49. Manly, F.J.B. Randomization, Bootstrap and Monte Carlo Methods in Biology, 2nd edn. London: Chapman and Hall/CRC (2001).

50. Excoffier, L. &Lischer, H.E.L. Arlequin suite ver 3.5: a new series of programs to perform population genetics analyses under Linux and Windows. Mol. Ecol. Resour. 10, 564–567 (2010).

51. Spackman, M.E. & McKechnie, S.W. Assessing the value of mitochondrial DNA variation for detecting population subdivision in the cotton bollworm, *Helicoverpa armigera* (Lepidoptera: Noctuidae), in Australia. Proc. Beltwide Cotton Conf. 2, 811–813 (1995).

52. Behere, G.T. et al. Mitochondrial DNA analysis of field populations of *Helicoverpa armigera* (Lepidoptera: Noctuidae) and of its relationship to *H. zea*. BMC Evol. Biol. 7, 117–127 (2007).

53. Daly, J.C. & Gregg, P. Genetic variation in *Heliothis* in Australia: species identification and gene flow in the two pest species *H. armigera* (Hübner) and *H. punctigera* Wallengren (Lepidoptera: Noctuidae). B. Entomol. Res. 75, 169–184 (1985).

54. Zhou, X., Faktor, O., Applebaum, S.W. & Coll, M. Population structure of the pestiferous moth *Helicoverpa armigera* in the eastern Mediterranean using RAPD analysis. Heredity (Edinb) 85, 251–256 (2000).

55. Xiao-feng, C., Sheng-jiang, T., Ren-yi, L., Ying, W. & Diamo, L. Study on the genetic variation of the cotton bollworm *Helicoverpa armigera* (Hübner) populations in china. Insect Sci. 7, 243–249 (2000).

56. Endersby, N.M., Hoffmann, A.A., McKechnie, S.W. & Weeks, A.R. Is there genetic structure in populations of *Helicoverpa armigera* from Australia? Entomol. Exp. Appl. 122, 253–263 (2007).

57. Vassal, J.M., Brevault, T., Achaleke, J. & Menozzi, P. Genetic structure of the polyphagous pest *Helicoverpa armigera* (Lepidoptera: Noctuidae) across the sub-Saharan cotton belt. Commun. Agric. Appl. Biol. Sci. 73, 433–437(2008).

58. Anderson, C.J., Tay, W.T., McGaughran, A., Gordon, K. & Walsh, T.K. Population structure and gene flow in the global pest *Helicoverpa armigera*. Mol. Ecol. 25, 5296–5311(2016).

59. Tay, W.T., Evans, G.A., Boykin, L.M. & De Barro, P.J. Will the real *Bemisia tabaci* please stand up? PLoS ONE 7(11), e50550; 10.1371/journal.pone.0050550 (2012)

60. Grille, G., Gauthier, N., Buenahora, J., Basso, C. & Bonato, O. First report of the Q biotype of *Bemisia tabaci* in Argentina and Uruguay. Phytoparasitica 39, 235–238 (2011).

61. de Moraes, L.A., Marubayashi, J.M., Yuki, V.A., Ghanim, M., Bello, V.H., De Marchi, B.R., Barbosa, L.D., Boykin, L.M., Krause-Sakate, R. & Pavan, M.A. New invasion of *Bemisia tabaci* Mediterranean species in Brazil associated to ornamental plants. Phytoparasitica 45, 517–525 (2017).

62. da Fonseca Barbosa, L., Yuki, V.A., Marubayashi, J.M., De Marchi, B.R., Perini, F.L., Pavan, M.A., de Barros, D.R., Ghanim, M., Moriones, E., Navas-Castillo, J. et al. First report of *Bemisia tabaci* Mediterranean (Q biotype) species in Brazil. Pest Management Science 71, 501–504 (2015).

63. Gaither, M.R., Bowen, B.W. & Toonen, R.J. Population structure in the native range predicts the spread of introduced marine species. P. Roy. Soc. B-Biol. Sci. 280, 20130409; 10.1098/rspb.2013.0409 (2013).

64. Oliveira, H.N., Simonato, J., Glaeser, D.F. & Pereira, F.F. Parasitism of *Helicoverpa armigera* pupae (Lepidoptera: Noctuidae) by *Tetrastichus howardi* and *Trichospinus diatraeae* (Hymenoptera: Eulophidae). Semina. Ciencias Agrárias 37, 111–115 (2016).

65. Lockwood, J.L., Casse, P. & Blackburn, T. The role of propagule pressure in explaining species invasions. Trends. Ecol. Evol. 20, 223–228 (2005).

66. Simberloff, D. The role of propagule pressure in biological invasions. Annu. Rev. Ecol. Evol. S. 40, 81–102 (2009).

67. Benjamini, Y. & Y. Hochberg. Controlling the false discovery rate: a practical and powerful approach to multiple testing. Journal of the Royal Statistical Society B 57, 289–300 (1995).

68. McDonald, J.H. 2014. Handbook of Biological Statistics, 3rd ed. Sparky House Publishing, Baltimore, Maryland.

69. Chown, S.L et al. Biological invasions, climate change and genomics. Evol. Appl. 8, 23–46 (2014).

70. Pearce, S.L. et al. Genomic innovations, transcriptional plasticity and gene loss underlying the evolution and divergence of two highly polyphagous and invasive *Helicoverpa* pest species. BMC Biol. 15, 69 (2017).

71. Mallet, J., Korman, A., Heckel, D.G. & King, P. Biochemical genetics of *Heliothis* and *Helicoverpa* (Lepidoptera, Noctuidae) and evidence for a founder event in *Helicoverpa zea*. Ann. Ent. Soc. Amer. 86, 189–197 (1993).

72. Guidolin, A.S., Fresia, P. & Cônsoli, F.L. The genetic structure of an invasive pest, the Asian citrus psyllid *Diaphorinacitri* (Hemiptera: Liviidae). PLoS ONE 9, e115749; 10.1371/journal.pone.0115749 (2014).

73. Walsh, T.K., Joussen, N., Tian, K., McGaughran, A., Anderson, C.J., Qiu. X. et al. Multiple recombination events between two cytochrome P450 loci contribute to global pyrethroid resistance in Helicoverpa armigera. PLoS ONE 13(11): e0197760. https://doi.org/10.1371/journal.pone.0197760 (2018).

74. Yang, Y., Chen, H., Wu, Y., Yang, Y. & Wu, S. Mutated cadherin alleles from a field population of *Helicoverpa armigera* confer resistance to *Bacillus thuringiensis* toxin Cry1Ac. Appl. Environ. Microb. 73, 6939–6944 (2007).

75. Achaleke, J. & Brevault, T. Inheritance and stability of pyrethroid resistance in the cotton bollworm *Helicoverpa armigera* (Lepidoptera: Noctuidae) in Central Africa. Pest Manag. Sci. 66, 137–141 (2010).

76. Zhang, H., Wu, S., Yang, Y., Tabashnik, B.E. & Wu, Y. Non-recessive Bt toxin resistance conferred by an intracellular cadherin mutation in field-selected populations of cotton bollworm. PLoS ONE 7, e53418; 10.1371/journal.pone.0053418 (2012).

77. Durigan, M.R. et al. High frequency of CYP337B3 gene associated with control failures of *Helicoverpa armigera* with pyrethroid insecticides in Brazil. Pesticide Biochemistry and Physiology; 10.1016/j.pestbp.2017.09.005 (2017).

78. Nair, R., Kalia, V., Aggarwal, K.K. & Gujar, G.T. Variation in the cadherin gene sequence of Cry1Ac susceptible and resistant *Helicoverpa armigera* (Lepidoptera: Noctuidae) and the identification of mutant alleles in resistant strains. Curr. Sci. 104, 215 (2013).

79. Wares, J.P., Hughes, A.R. & Grosberg, R.K. Mechanisms that drive evolutionary change: insights from species introductions and invasions in Species invasions: Insights into ecology, evolution, and biogeography (ed. Sax, D.F., Stachowicz, J.J., Gaines, S.D.) 229–257. Massachusetts: Sinauer Associates (2005).

80. Ellestrand, N.C. & Schierenbeck, K.A. Hybridization as a stimulus for the evolution of invasiveness in plants? Proc. Natl. Acad. Sci. U S A97, 7043–7050 (2000).

81. De Barro, P.J., Liu, S.S., Boykin, L.M. & Dinsdale, A.B. *Bemisia tabaci*: a statement of species status. Annu Rev Entomol 56, 1–19 (2011).

