## Supplementary material for "Multiple incursion pathways for *Helicoverpa armigera* in Brazil show its genetic diversity spreading in a connected world": Supp info

**Suppl. Table 1:** South American and global *Helicoverpa armigera* sequences and iBoL/GenBank accession numbers for the mtDNA COI 5’ region. Numbers of individuals (n) from each location are indicated in parentheses. Haplotypes identified in this study are indicated by †

| **Countries** | **Locations (n)** | **iBoL/GenBank accession numbers** |
| --- | --- | --- |
| **China** | Kunming (8) | GQ892840.1, GQ892842.1, GQ892854.1, GQ995232.1, GQ995234.1, GQ995235.1, GQ995244.1, GQ995239.1 |
|  | Tibet (2) | JX392497.1, JX392415.1 |
|  | Miaofengshan (4) | JX509766.1, JX509765.1, JX509764.1, JX509739.1 |
|  | Not specified (1) | HQ132369.1 |
|  | Dali (15) | GQ892846.1, GQ892847.1, GQ892848.1, GQ892849.1, GQ892850.1, GQ892851.1, GQ892852.1, GQ892853.1, GQ995233.1, GQ995236.1, GQ995237.1, GQ995240.1, GQ995241.1, GQ995242.1, GQ995243.1 |
|  | Henan (2) | GQ995238.1, GQ892855.1 |
|  | Lijiang (1) | GQ892845.1 |
|  | Yuxi (2) | GQ892843.1, GQ892844.1 |
| **Thailand** | Not specified (1) | EU768935.1 |
| **India** | Not specified (6) | HM854928.1, HM854929.1, HM854930.1, HM854931.1, HM854932.1, JX532104.1 |
| **Pakistan** | Not specified (2) | JN988529.1, JN988530.1 |
| **Europe** | Not specified (24) | FN907979.1, FN907980.1, FN907988.1, FN907989.1, FN907995.1, FN907996.1, FN907997.1, FN907998.1, FN907999.1, FN908000.1, FN908001.1, FN908002.1, FN908003.1, FN908005.1, FN908006.1, FN908011.1, FN908013.1, FN908014.1, FN908015.1, FN908016.1, FN908017.1, FN908018.1, FN908023.1, FN908026.1 |
|  | Germany (4) | GU654969.1, GU686757.1, GU686955.1, JF415782.1 |
| **Australia** | Queensland (10) | ANICL283-10, ANICL284-10, GBGL12943-14, GBGL12944-14, GBGL12945-14, GBGL12946-14, GBGL12947-14, GBGL12948-14, GBGL12949-14, GBGL12950-14 |
|  | New South Wales (8) | GBGL12935-14, GBGL12936-14, GBGL12937-14, GBGL12938-14, GBGL12939-14, GBGL12940-14, GBGL12941-14, GBGL12942-14 |
| **Brazil** | Bahia (112) | KM274936-KM274938, KM27513-KM275140 KM274939-KM274941, KM274943-KM274950, KM274951-KM274953, KM274957-KM274975, KM274979-KM274986, KM275038-KM275052, KM275078-KM275082, KM275070-KM275077, KM275127-KM275136, KF624850-KF624861 |
|  | Maranhão (20) | KM27498-KM274996, KM275103-KM275112 |
|  | Mato Grosso (19) | KM275083-KM275092, KM275156-KM275158, KM275097-KM275102 |
|  | Piaui (39) | KF624811- KF624849 |
|  | Roraima (14) | KF624862-KF624875 |
|  | Parana (3) | † (MG230503 - MG230505) |
|  | Santa Catarina (7) | † (MG230518 - MG230524) |
|  | Rio Grande do Sul (12) | † (MG230506 - MG230517) |
| **Argentina** | Santa Fé (3) | † (MG230495 - MG230497) |
| **Uruguay** | Maldonado (2) | † (MG230525 - MG230526 (and sequences from Arnemann et al [7]) |
| **Paraguay** | Alto Paraná (10) | † (MG230499 - MG230502  (and sequences from Arnemann et al [7]) |

**Suppl. Table 2:** Detected mtDNA COI haplotypes within South America locations with the number of *Helicoverpa armigera* individuals carrying specific haplotypes at particular sites indicated.

| **Haplotypes** | **ARG** | **URY** | **PRY** | **Brazilian states** | | | | | | | |  | |
| --- | --- | --- | --- | --- | --- | --- | --- | --- | --- | --- | --- | --- | --- |
|  |  |  |  | **BA** | **MA** | **MT** | **PI** | **RR** | **PR** | **SC** | **RS** |  | **n** |
| **Harm_BC01** | 0 | 0 | 3 | 24 | 8 | 6 | 11 | 6 | 1 | 4 | 3 |  | 66 |
| **Harm_BC02** | 0 | 1 | 0 | 36 | 9 | 6 | 18 | 5 | 1 | 2 | 3 |  | 81 |
| **Harm_BC03** | 1 | 0 | 0 | 13 | 1 | 6 | 3 | 1 | 0 | 0 | 0 |  | 25 |
| **Harm_BC04** | 1 | 0 | 0 | 10 | 0 | 0 | 1 | 0 | 0 | 0 | 4 |  | 16 |
| **Harm_BC05** | 0 | 0 | 0 | 5 | 0 | 0 | 2 | 1 | 1 | 0 | 0 |  | 9 |
| **Harm_BC06** | 0 | 0 | 0 | 0 | 1 | 0 | 0 | 0 | 0 | 0 | 0 |  | 1 |
| **Harm_BC07** | 0 | 0 | 0 | 2 | 0 | 0 | 3 | 0 | 0 | 0 | 0 |  | 5 |
| **Harm_BC13** | 0 | 0 | 2 | 0 | 0 | 0 | 0 | 0 | 0 | 1 | 0 |  | 3 |
| **Harm_BC14** | 0 | 0 | 0 | 2 | 0 | 0 | 0 | 0 | 0 | 0 | 0 |  | 2 |
| **Harm_BC16** | 1 | 0 | 1 | 0 | 0 | 0 | 0 | 0 | 0 | 0 | 0 |  | 2 |
| **Harm_BC17** | 0 | 0 | 2 | 0 | 0 | 0 | 0 | 0 | 0 | 0 | 0 |  | 2 |
| **Harm_BC23** | 0 | 0 | 0 | 0 | 0 | 0 | 1 | 0 | 0 | 0 | 0 |  | 1 |
| **Harm_BC24** | 0 | 0 | 0 | 0 | 0 | 0 | 0 | 1 | 0 | 0 | 0 |  | 1 |
| **Harm_BC34** | 0 | 0 | 0 | 1 | 0 | 0 | 0 | 0 | 0 | 0 | 0 |  | 1 |
| **Harm_BC35** | 0 | 0 | 0 | 1 | 0 | 0 | 0 | 0 | 0 | 0 | 0 |  | 1 |
| **Harm_BC36** | 0 | 0 | 0 | 1 | 0 | 0 | 0 | 0 | 0 | 0 | 0 |  | 1 |
| **Harm_BC37** | 0 | 0 | 0 | 0 | 1 | 0 | 0 | 0 | 0 | 0 | 0 |  | 1 |
| **Harm_BC38** | 0 | 0 | 0 | 0 | 0 | 1 | 0 | 0 | 0 | 0 | 0 |  | 1 |
| **Harm_BC39** | 0 | 0 | 0 | 1 | 0 | 0 | 0 | 0 | 0 | 0 | 0 |  | 1 |
| **Harm_BC42** | 0 | 0 | 0 | 0 | 0 | 0 | 0 | 0 | 0 | 0 | 1 |  | 1 |
| **Harm_BC43** | 0 | 0 | 1 | 0 | 0 | 0 | 0 | 0 | 0 | 0 | 0 |  | 1 |
| **Harm_BC44** | 0 | 1 | 0 | 0 | 0 | 0 | 0 | 0 | 0 | 0 | 0 |  | 1 |
| **Harm_BC45** | 0 | 0 | 0 | 0 | 0 | 0 | 0 | 0 | 0 | 0 | 1 |  | 1 |
| **Harm_BC46** | 0 | 0 | 1 | 0 | 0 | 0 | 0 | 0 | 0 | 0 | 0 |  | 1 |
| **Harm_BC47** | 1 | 0 | 0 | 0 | 0 | 0 | 0 | 0 | 0 | 0 | 0 |  | 1 |
| **n** | 4 | 2 | 10 | 96 | 20 | 19 | 39 | 14 | 3 | 7 | 12 |  | 226 |

**Suppl. Table 3:** Haplotypes distribution within individuals from Australia (AUS, n = 18), Asia (Asia, n = 44), Europe (EUR, n = 28), Argentina (ARG, n = 4), Uruguay (URY, n = 2), Paraguay (PRY, n = 10) and Brazilian states (Bahia BA, n = 96; Maranhão MA, n = 20; Mato Grosso MT, n = 19; Piauí PI, n = 39; Roraima RR, n = 14; Paraná PR, n = 3; Santa Catarina SC, n = 7; Rio Grande do Sul RS, n = 12).

| **Haplotypes** | **AUS** | **ASIA** | **EUR** | **ARG** | **URY** | **PRY** | **BRAZILIAN STATES** | | | | | | | |
| --- | --- | --- | --- | --- | --- | --- | --- | --- | --- | --- | --- | --- | --- | --- |
|  |  |  |  |  |  |  | **BA** | **MA** | **MT** | **PI** | **RR** | **PR** | **SC** | **RS** |
| **Harm_BC01** | 2 | 10 | 14 |  |  | 3 | 24 | 8 | 6 | 11 | 6 | 1 | 4 | 3 |
| **Harm_BC02** |  | 4 | 2 |  | 1 |  | 36 | 9 | 6 | 18 | 5 | 1 | 2 | 3 |
| **Harm_BC03** | 1 |  | 3 | 1 |  |  | 13 | 1 | 6 | 3 | 1 |  |  |  |
| **Harm_BC04** |  | 3 | 3 | 1 |  |  | 10 |  |  | 1 |  |  |  | 4 |
| **Harm_BC05** |  |  |  |  |  |  | 5 |  |  | 2 | 1 | 1 |  |  |
| **Harm_BC06** |  | 6 | 2 |  |  |  |  | 1 |  |  |  |  |  |  |
| **Harm_BC07** |  | 1 |  |  |  |  | 2 |  |  | 3 |  |  |  |  |
| **Harm_BC08** | 5 |  |  |  |  |  |  |  |  |  |  |  |  |  |
| **Harm_BC09** |  | 5 |  |  |  |  |  |  |  |  |  |  |  |  |
| **Harm_BC10** |  | 3 | 1 |  |  |  |  |  |  |  |  |  |  |  |
| **Harm_BC11** | 3 |  |  |  |  |  |  |  |  |  |  |  |  |  |
| **Harm_BC12** | 3 |  |  |  |  |  |  |  |  |  |  |  |  |  |
| **Harm_BC13** |  |  |  |  |  | 2 |  |  |  |  |  |  | 1 |  |
| **Harm_BC14** |  |  |  |  |  |  | 2 |  |  |  |  |  |  |  |
| **Harm_BC15** |  | 2 |  |  |  |  |  |  |  |  |  |  |  |  |
| **Harm_BC16** |  |  |  | 1 |  | 1 |  |  |  |  |  |  |  |  |
| **Harm_BC17** |  |  |  |  |  | 2 |  |  |  |  |  |  |  |  |
| **Harm_BC18** | 2 |  |  |  |  |  |  |  |  |  |  |  |  |  |
| **Harm_BC19** |  | 1 |  |  |  |  |  |  |  |  |  |  |  |  |
| **Harm_BC20** |  |  | 1 |  |  |  |  |  |  |  |  |  |  |  |
| **Harm_BC21** |  |  | 1 |  |  |  |  |  |  |  |  |  |  |  |
| **Harm_BC22** |  |  | 1 |  |  |  |  |  |  |  |  |  |  |  |
| **Harm_BC23** |  |  |  |  |  |  |  |  |  | 1 |  |  |  |  |
| **Harm_BC24** |  |  |  |  |  |  |  |  |  |  | 1 |  |  |  |
| **Harm_BC25** |  | 1 |  |  |  |  |  |  |  |  |  |  |  |  |
| **Harm_BC26** |  | 1 |  |  |  |  |  |  |  |  |  |  |  |  |
| **Harm_BC27** |  | 1 |  |  |  |  |  |  |  |  |  |  |  |  |
| **Harm_BC28** |  | 1 |  |  |  |  |  |  |  |  |  |  |  |  |
| **Harm_BC29** |  | 1 |  |  |  |  |  |  |  |  |  |  |  |  |
| **Harm_BC30** |  | 1 |  |  |  |  |  |  |  |  |  |  |  |  |
| **Harm_BC31** |  | 1 |  |  |  |  |  |  |  |  |  |  |  |  |
| **Harm_BC32** |  | 1 |  |  |  |  |  |  |  |  |  |  |  |  |
| **Harm_BC33** |  | 1 |  |  |  |  |  |  |  |  |  |  |  |  |
| **Harm_BC34** |  |  |  |  |  |  | 1 |  |  |  |  |  |  |  |
| **Harm_BC35** |  |  |  |  |  |  | 1 |  |  |  |  |  |  |  |
| **Harm_BC36** |  |  |  |  |  |  | 1 |  |  |  |  |  |  |  |
| **Harm_BC37** |  |  |  |  |  |  |  | 1 |  |  |  |  |  |  |
| **Harm_BC38** |  |  |  |  |  |  |  |  | 1 |  |  |  |  |  |
| **Harm_BC39** |  |  |  |  |  |  | 1 |  |  |  |  |  |  |  |
| **Harm_BC40** | 1 |  |  |  |  |  |  |  |  |  |  |  |  |  |
| **Harm_BC41** | 1 |  |  |  |  |  |  |  |  |  |  |  |  |  |
| **Harm_BC42** |  |  |  |  |  |  |  |  |  |  |  |  |  | 1 |
| **Harm_BC43** |  |  |  |  |  | 1 |  |  |  |  |  |  |  |  |
| **Harm_BC44** |  |  |  |  | 1 |  |  |  |  |  |  |  |  |  |
| **Harm_BC45** |  |  |  |  |  |  |  |  |  |  |  |  |  | 1 |
| **Harm_BC46** |  |  |  |  |  | 1 |  |  |  |  |  |  |  |  |
| **Harm_BC47** |  |  |  | 1 |  |  |  |  |  |  |  |  |  |  |

**Suppl. Table 4:** Observations of haplotypes within South American locations summed into two broad categories.

| **Haplotypes** | **Non-Brazil** | **Brazil** | **n** |
| --- | --- | --- | --- |
| Harm_BC01 | 3 | 63 | 66 |
| Harm_BC02 | 1 | 80 | 81 |
| Harm_BC03 | 1 | 24 | 25 |
| Harm_BC04 | 1 | 15 | 16 |
| Harm_BC05 | 0 | 9 | 9 |
| Harm_BC06 | 0 | 1 | 1 |
| Harm_BC07 | 0 | 5 | 5 |
| Harm_BC13 | 2 | 1 | 3 |
| Harm_BC14 | 0 | 2 | 2 |
| Harm_BC16 | 2 | 0 | 2 |
| Harm_BC17 | 2 | 0 | 2 |
| Harm_BC23 | 0 | 1 | 1 |
| Harm_BC24 | 0 | 1 | 1 |
| Harm_BC34 | 0 | 1 | 1 |
| Harm_BC35 | 0 | 1 | 1 |
| Harm_BC36 | 0 | 1 | 1 |
| Harm_BC37 | 0 | 1 | 1 |
| Harm_BC38 | 0 | 1 | 1 |
| Harm_BC39 | 0 | 1 | 1 |
| Harm_BC42 | 0 | 1 | 1 |
| Harm_BC43 | 1 | 0 | 1 |
| Harm_BC44 | 1 | 0 | 1 |
| Harm_BC45 | 0 | 1 | 1 |
| Harm_BC46 | 1 | 0 | 1 |
| Harm_BC47 | 1 | 0 | 1 |
| n | 16 | 210 | 226 |

**Suppl. Table 5:** Pairwise *F*_ST_ estimates of 11 populations of *Helicoverpa armigera* from Brazilian states, Argentina, Uruguay and Paraguay.

|  | **Roraima** | **Maranhão** | **Piauí** | **Bahia** | **Mato Grosso** | **Santa Catarina** | **Rio Grande do Sul** | **Paraná** | **Argentina** | | | **Uruguai** | **Paraguay** |
| --- | --- | --- | --- | --- | --- | --- | --- | --- | --- | --- | --- | --- | --- |
| **Roraima** | - |  |  |  |  |  |  |  |  | | |  |  |
| **Maranhão** | -0.0503 | - |  |  |  |  |  |  |  | | |  |  |
| **Piauí** | -0.0313 | -0.0046 | - |  |  |  |  |  |  | | |  |  |
| **Bahia** | -0.0199 | -0.0088 | -0.0001 | - |  |  |  |  |  | | |  |  |
| **Mato Grosso** | 0.0121 | 0.0111 | 0.0378 | 0.0179 | - |  |  |  |  | | |  |  |
| **Santa Catarina** | 0.0160 | 0.0312 | 0.0977 | 0.0506 | 0.1181 | - |  |  |  | | |  |  |
| **Rio Grande do Sul** | 0.0615 | 0.0862 | **0.1203** | **0.0630** | **0.1327** | -0.0096 | - |  |  | | |  |  |
| **Paraná** | -0.0800 | 0.0395 | -0.0643 | -0.0316 | 0.0669 | 0.1102 | 0.0314 |  | |  | |  |  |
| **Argentina** | 0.1917 | **0.2520** | **0.2661** | **0.1838** | 0.1324 | 0.1433 | 0.0518 | 0.0429 | - | |  | |  |
| **Uruguai** | 0.1152 | 0.2261 | 0.1009 | 0.0836 | 0.1310 | 0.3243 | 0.0400 | -0.0714 | -0.1429 | | | - |  |
| **Paraguay** | **0.3470** | **0.3947** | **0.4266** | **0.3759** | **0.3813** | **0.1832** | **0.1922** | **0.3010** | 0.1033 | | | 0.2214 | - |

**Note:** Uruguay sample size of *H. armigera* is n=2, of which one is the most common Harm_BC02 mtDNA COI haplotype detected in Brazil in this study, and the other being a rare mtDNA COI haplotype Harm_BC44 (see Suppl. Fig. 2). Sample size for Argentina is n=4, of which two haplotypes (i.e., Harm_BC03, Harm_BC04) representing the 3^rd^ and 4^th^ most commonly detected haplotypes were shared with Brazilian populations, one haplotype shared with Paraguay (i.e., Harm_BC16) and one unique haplotype (Harm_BC47). Of the n=10 *H. armigera* sampled in Paraguay, non-Brazilian haplotypes were detected in n=6, one rare haplotype (Harm_BC13) was shared with *H. armigera* from the state of Santa Catarina, and the remaining n=3 have a common Harm_BC01 haplotype found widely in Brazil. Of the n=12 *H. armigera* sampled from the State of Rio Grande do Sul, two unique haplotypes were detected (i.e., Harm_BC42, Harm_BC45).

**Suppl. Table 6:** Pairwise *F*_st_ estimates between Brazilian, Old World, and non-Brazilian (Argentina/Paraguay/Uruguay) *Helicoverpa armigera* populations.

| Population | Non-Brazilian | Brazilian |
| --- | --- | --- |
| Brazilian | 0.2879 | - |
| Old World | 0.2022 | 0.0742 |

**Suppl. Table 7:** Summary of AMOVA of different hierarchical population structures of *Helicoverpa armigera* from the South American continent. *p* < 0.05 indicated by ‘*’.

| Source of variation | d.f. | Sum of squares | Variance components | Percentage of variation | F-Statistic |
| --- | --- | --- | --- | --- | --- |
| Among groups | 3 | 10.219 | 0.04905 va | 7.10 | 0.07102* |
| Among populations within groups | 7 | 6.219 | 0.01919 vb | 2.78 | 0.02992 |
| Within populations | 215 | 133.814 | 0.62239 vc | 90.12 | 0.09881* |
| Total | 225 | 150.252 | 0.69063 |  |  |

**Suppl. Fig. 1**: Distribution of P-values from 10,000 significance tests using pseudo-observed data conforming to the null hypothesis. (a) Model validation results for matrices of size 25 rows x 11 columns, corresponding to the dimensions of the full dataset (Suppl. Table 3). (b) Model validation results for matrices of size 25 rows x 2 columns, corresponding to the dimensions of the aggregated dataset for examining patterns between Brazil and non-Brazilian locations (see above).

(a)	(b)

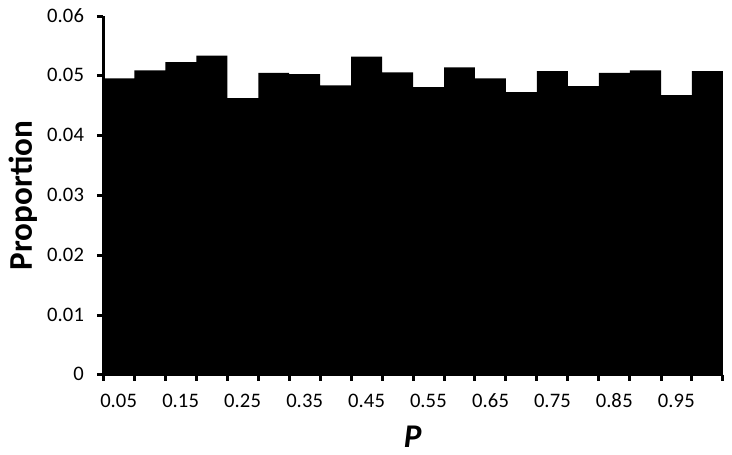

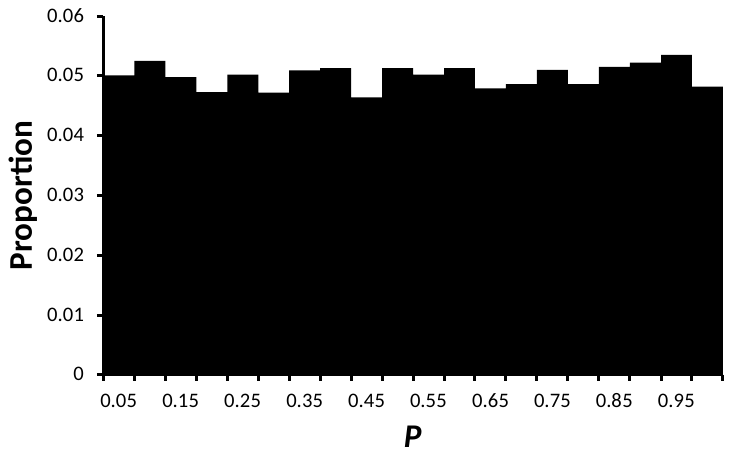

**Suppl. Fig. 2:** Haplotype distribution patterns and diversity of *Helicoverpa armigera* in South America based on the partial mtDNA COI gene (548bp). Pie charts (i), (ii) and (iii) are from Mato Grosso, Piaui and Bahia States, respectively.

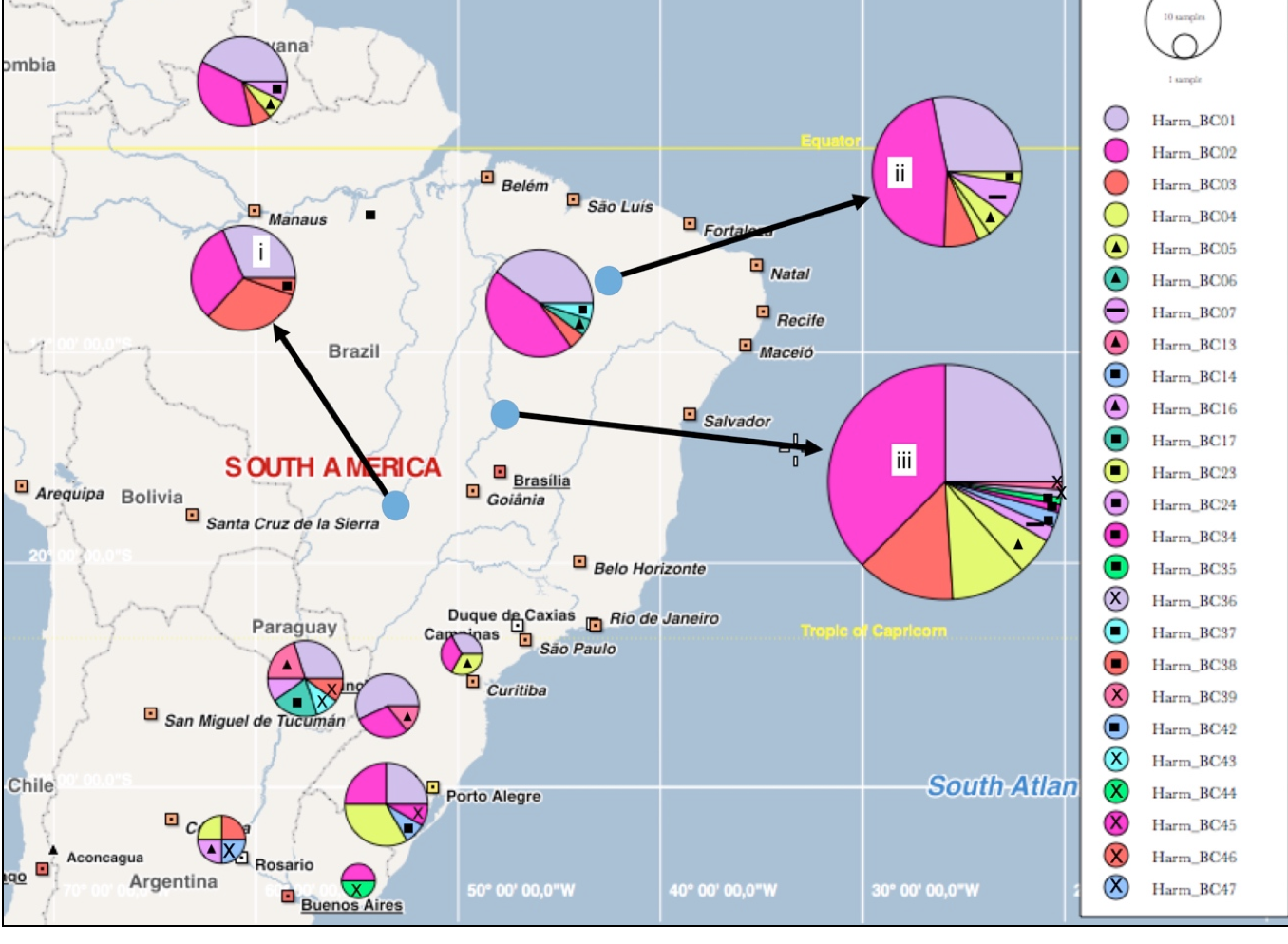

Generated using PopART <http://popart.otago.ac.nz>. Leigh, JW, Bryant D (2015). PopART: Full-feature software for haplotype network construction. Methods Ecol Evol 6(9):1110–1116.
